# Community composition drives metabolic competition and *Staphylococcus aureus* colonization resistance in synthetic nasal communities

**DOI:** 10.64898/2026.01.08.698450

**Authors:** Marcelo Navarro Diaz, Kerrin Bertram, Laura Camus, Soyoung Ham, Lars Angenent, Jose Antonio Velazquez Gomez, Simon Heilbronner, Paolo Stincone, Johanna Rapp, Daniel Petras, Hannes Link

## Abstract

The human nasal microbiome is a low-diversity ecosystem whose assembly principles and mechanisms of colonization resistance remain poorly understood. *Staphylococcus aureus* is a member of the nasal microbiome of some individuals with variable abundance. We hypothesized that nutritional competition, strain-level diversity, and nutrient availability shape community stability and the ability of commensal species to inhibit *S. aureus*. To test this, we constructed 50 defined synthetic communities composed of representative human nasal bacteria differing in strain and species composition and tracked their temporal dynamics, *S. aureus* growth, metabolic profiles, and nutritional interactions. The composition of synthetic communities with 5-10 species showed robust and reproducible dynamics and converged in one of three stable states. Synthetic communities dominated by a specific strain of *Corynebacterium propinquum* were highly stable and consistently excluded *S. aureus*. Growth curves and coculture assays showed that *C. propinquum* outcompetes *S. aureus* under poor nutritional conditions resembling the nasal environment, whereas *S. aureus* dominates in nutrient-rich conditions. Metabolomics analyses revealed that nutritional competition, including siderophores utilization and amino acid limitation, likely underlies this colonization resistance. These results establish a tractable synthetic community model for the human nasal microbiome and identify nutrient-dependent competition and microbial metabolite production as key drivers of community structure and pathogen exclusion.

## Introduction

The human microbiome is important for human health and provides protection against pathogens. This colonization resistance can occur through various mechanisms such as nutritional competition or by use of inhibitory compounds [1]. Each anatomical site contains distinct microbial communities composed of commensals, pathobionts, and pathogens, which may be permanent residents or transient colonizers. Among these sites, the nasal cavity is of particular interest because it is an entry point of airborne pathogens such as *Staphylococcus aureus*, which is a major human pathobiont [2]. Approximately one-quarter of adults are persistently colonized with *S. aureus* in the anterior nares, while others are intermittently colonized or remain uncolonized [3]. In *S. aureus* carriers, the nasal cavity functions as a reservoir from which *S. aureus* can disseminate to other body sites, occasionally leading to disease [4], which is especially dangerous in immunocompromised and post-operation patients [5, 6].

Despite its clinical importance, the ecological and mechanistic factors that determine the presence, absence, or abundance of *S. aureus* in the nasal microbiome are not completely understood. Colonization resistance to *S. aureus* in the nose is a complex process where the host’s immunity has been shown to play a role [7], and several species present in the nasal microbiome were shown to exhibit different types of interactions with *S. aureus* [8]. Species reported to inhibit the growth of *S. aureus* include *Staphylococcus lugdunensis* (which produces the antimicrobial lugdunin), *Staphylococcus epidermidis* [8], and *Dolosigranulum pigrum* [9]. Other important mechanisms of *S. aureus* exclusion, besides antimicrobial production, include nutrient depletion [10, 11] and siderophore-mediated competition [12]. Such nutritional competition is especially relevant in the context of the nasal environment, which has been shown to be limited in carbon and energy sources, nutrients and iron, shaping microbial interactions and colonization dynamics [10, 13]. However, disentangling these mechanisms *in vivo* and in the context of a whole microbial community is challenging, as the nasal cavity is a highly heterogeneous and host-specific environment that supports a comparatively small, but highly specialized, microbial community [14, 15].

By using synthetic communities (SynComs), the low-diversity nasal microbiome provides an ideal model to study fundamental ecological processes, such as community assembly, metabolic cross-feeding, and invasion dynamics. SynComs have been widely used to model other microbiome systems, such as the human gut microbiome [16, 17] and the rhizosphere of plants [18, 19]. These model systems have been instrumental in dissecting mechanisms of community assembly [18, 20], microbial function and metabolic interactions [21, 22], and responses to perturbations such as antibiotics or pathogen invasion [19, 23]. Previously, the PNC8 SynCom was developed as a model of the piglet nasal microbiome [24]. In contrast, to our knowledge there no SynComs for the human nasal microbiome, only pairwise pairwise cultures assembled from intrapatient strains [25]. Thus, an experimentally tractable SynCom system to study metabolic interactions and colonization resistance against *Staphylococcus aureus* in the human nasal ecosystem is currently lacking.

Seven community state types (CSTs) have been described for the human nasal microbiome based on species-level composition of *in vivo* datasets, suggesting recurring patterns of co-occurrence among dominant taxa [26]. However, these CSTs were defined on species-level resolution and do not account for strain-level variation, which may substantially influence ecological interactions and assembly dynamics. Whether nasal CSTs represent deterministic ecological attractors emerging from intrinsic microbial interactions, or instead reflect host filtering and historical contingency, remains unclear. To address this question, we adopted an unbiased assembly strategy using prevalent nasal species combined randomly to test whether stable community structures resembling in vivo CSTs would emerge from intrinsic interaction dynamics and how such configurations influence resistance to *S. aureus* invasion.

Here, we describe the development of a set of SynComs using a previously reported culture collection of human nasal isolates [25]. We assembled 50 SynComs containing random combinations of the most prevalent nasal bacterial species, each represented by three distinct strains, and cultured them on agar plates containing a synthetic nasal medium, previously developed to resemble the nutritional conditions of the nose [10]. After SynComs were established, we added *S. aureus* to test their colonization resistance. We characterized their compositional and metabolic profiles using long-read amplicon sequencing and untargeted and targeted metabolomics. Our approach allowed us to identify key species, strain-level effects, and metabolite signatures that are associated with abundance of *S. aureus* in the SynComs. These results provide a tractable *in vitro* system to explore the ecological determinants of nasal community assembly and pathogen exclusion.

## Methods

### Bacterial pool selection

To select species for SynCom construction, we first analyzed the Human Microbiome Project (HMP) 16S rRNA amplicon sequencing data from the *nasal cavity* body site [27]. Raw sequences were processed using QIIME 2 v2021.11 [28]. Quality filtering, denoising, and chimera removal were performed with DADA2 [29], followed by inference of amplicon sequence variants (ASVs). Taxonomic assignment was carried out using BLAST [30] against the NCBI 16S rRNA RefSeq database [31]. The complete ASVs table is provided in Supplementary Table S1.

Experiments were performed using bacterial isolates from the LaCa collection of nasal bacterial isolates [25]. The LaCa collection comprises 228 isolates of bacteria (119 unique strains of 33 different species) from the nasal cavities of 11 healthy volunteers [25]. We selected 10 of the most prevalent species based on our HMP analysis (Supplementary Figure S1), with the limitation that they were available in the LaCa collection and selecting 3 strains per species isolated from different volunteers to ensure that isolates did not originate from the same population. For two species, for which the LaCa collection did not have enough interpatient strains (*Corynebacterium pseudodiphtheriticum* and *Corynebacterium tuberculostearicum*), we obtained 1 strain from the German Collection of Microorganisms and Cell Cultures (DSMZ). These were type strains *C. pseudodiphtheriticum* DSM44287 and *C. tuberculostearicum* DSM44922.

### SynCom cultivation

To avoid imposing predefined community structures, combinations were generated randomly rather than replicating previously described CST compositions. Fifty synthetic communities (SynComs) were assembled by randomly selecting both the number (from 5 to 10) and identity of species, along with one randomly chosen strain per species. This design enabled assessment of whether stable and recurrent community configurations would emerge from intrinsic interaction dynamics across strain backgrounds, thereby allowing evaluation of determinism versus contingency in nasal community assembly. The experimental workflow is depicted in (Figure 1), and species/strains used in each SynCom are listed in Supplementary Table S2.

**Figure 1.**
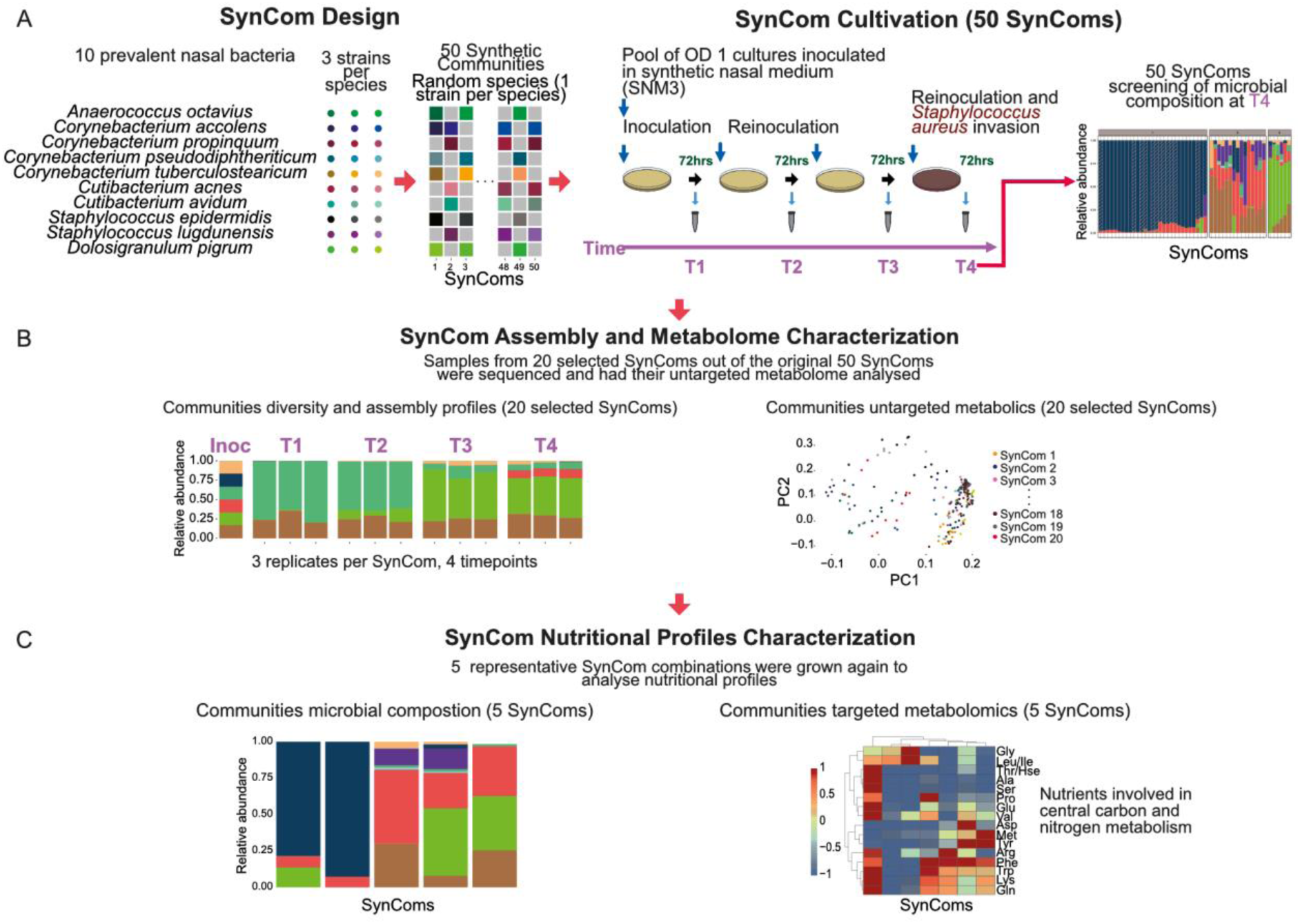
SynCom design and experimental workflow. (A) Ten of the most prevalent nasal bacterial species, each represented by three strains, were selected based on analyses of Human Microbiome Project data and the strain availability in the LaCa collection of nasal isolates [25]. SynComs were assembled by randomly selecting both the number (from 5 to 10) and identity of species per community, followed by the random selection of one strain per species. Communities were cultivated on Synthetic Nasal Medium agar plates (SNM3;[10]). SynComs were grown for 72 h, after which biomass was collected for reinoculation, DNA extraction and sequencing, and metabolomics analyses. Three sequential reinoculations on fresh SNM3 plates were performed. In the final reinoculation, *S. aureus* was introduced to assess colonization resistance. SynCom composition was determined using Nanopore sequencing. (B) Twenty selected communities were further characterized across all time points and replicates using sequencing and untargeted metabolomics to investigate assembly dynamics and metabolome composition. (C) Finally, five representative SynCom combinations were grown again from scratch to examine nutrient production and consumption profiles using targeted metabolomics.

Starter cultures were grown from glycerol stocks 24–72 h before experiments using BHI, LB, or Columbia agar plates, depending on species (Supplementary Table S3). Bacterial biomass was harvested from plates with sterile swabs, resuspended in PBS, and adjusted to OD₆₀₀ 1. Equal volumes (100 µL) of each strain suspension were pooled to generate the SynCom inoculum. SynComs were plated in triplicate on Synthetic Nasal Medium 3 (SNM3) agar plates, designed to mimic human nasal secretions [10], and incubated for 72 h at 34°C. Inoculation of pooled cultures was made on three SNM3 plates to obtain three technical replicates per SynCom. Biomass was collected with sterile loops using a star pattern across the plate. This pattern allowed us to collect enough biomass for DNA extraction while preserving enough biomass for metabolomics analyses. Biomass was resuspended in 500 µL PBS and used for reinoculation and DNA extraction. The rest of the biomass on the plate was collected for metabolomics by transferring it into tubes containing 1mL of 80% methanol and storing them at −80 °C until processing. Reinoculation consisted of diluting the PBS suspension to OD₆₀₀ 1 and plating 100 µL on fresh SNM3 plates. This process was repeated for three sequential transfers. In the final transfer, SynComs were challenged by inoculating *S. aureus* USA300 together with the SynCom reinoculate at a 1000-fold lower OD. Staphylococcus aureus USA300 is a dominant community-associated methicillin-resistant S. aureus (MRSA) clone that has emerged as a leading cause of severe infections worldwide [32].

### Repetition experiment

To evaluate reproducibility and analyze metabolites consumption profiles, five SynComs were cultivated in a separate experiment using 12-well plates containing 2 mL of SNM3 agar per well. Culture initiation and sampling followed the same procedure as in the main experiment, except that 20 µL of inoculum was used per well. Biomass was collected from the wells for DNA extraction and reinoculation. For targeted metabolomics, biomass was extracted by flooding from the wells with 2mL of 80% methanol, incubated at −20 °C for 20 min, and processed as described above. Wells without inoculum containing only SNM3 were analyzed as control. Additionally, *S. aureus* USA300 and the three strains of *C. propinquum* were also grown as monocultures in 12-well plates to analyze their metabolome. Three biological replicates were used for each SynCom, control and monocultures.

### Co-cultures

To test how the nutritional environment affected the interaction between *C. propinquum* and *S. aureus*, we performed co-cultures between the 3 strains of *C. propinquum* used in the SynCom experiments and *S. aureus* USA300. For these co-cultures we used SNM3, SNM10 (which contained 3x or 10x more amino acids concentration of base SNM), and the nutrient-rich medium BHI. We followed the same inoculation protocol of the SynCom experiment after which we took biomass samples for DNA extraction and sequencing. Three biological replicates were used for each of the co-cultures.

### Sequencing analyses

DNA was extracted using the ZymoBIOMICS DNA Miniprep Kit (Zymo Research Corp., Irvine, CA, USA) following the manufacturer’s protocol. Full-length rRNA operon amplicons (∼4,000 bp) were generated by PCR using primers 16S-27F-GGTGCTG-ONT and 23S-2241R-GGTGCTG-ONT as described previously [33]. Nanopore sequencing libraries were prepared with the Rapid Barcoding Kit (SQK-RBK114.96) following the “Amplicon sequencing from DNA protocol” (available at https://nanoporetech.com/document/rapid-sequencing-v14-amplicon-sequencing-sqk-rbk114-24-or-sqk) and sequenced on MinION V14 flow cells (Oxford Nanopore Technologies) using MinKNOW v25.05.14. Only reads with a quality score ≥10 were retained and demultiplexed. Taxonomic classification was performed using Emu v3.4.5 [34] with a custom rRNA operon reference database generated from the strain genomes via in silico PCR using EMBOSS-primersearch [35] as previously reported [33].

Initial screening involved sequencing one replicate of the final time point (*i.e.*, after *S. aureus* invasion) for each of the 50 SynComs. The complete table is provided in Supplementary Table S4. Twenty representative SynComs were then selected for sequencing of all time points and replicates. The complete table is provided in Supplementary Table S5. Tables for the repetition experiment and coculture experiments are available as Supplementary Table S6 and Supplementary Table S7 respectively. Raw fastq files from sequencing data are available in the NCBI SRA database under the BioProject PRJNA1370791.

### Growth curves

To analyze the differences in the growth of the species and their strains we performed growth curves in SNM3. Biomass from reactivated strains was transferred using an inoculation loop to BHI medium in a 5 ml culture tube and incubated overnight with shaking at 220 rpm and at 37°C. Turbidity was assessed to ensure sufficient growth before transferring 1.8 ml of medium to a sterile 2 ml Eppendorf tube. Cells were then washed twice in SNM salts base (SNM without glucose or amino acids). The OD was adjusted to 0.02 in SNM3 and cultures were transferred to a flat bottom 96-well plates. Cultures were incubated at 37°C and shaken continuously at 269 cpm (6 mm). The OD was measured every 10 min for 24 h using a BioTek SYNERGY H1 microplate reader (Agilnet, Santa Clara, CA, USA). The complete tables for growth curve data is provided in Supplementary Table S8 and Supplementary Table S9.

### Untargeted metabolomics

For the samples of 20 selected SynComs, we sonicated the biomass suspended in 80% methanol for five cycles (60s on, 10 s off, 40% amplitude), centrifuged (10,000 rpm, 10 min), and supernatants were transferred to pre-weighed HPLC glass vials. Methanol was evaporated under vacuum, and dried extracts were weighed and resuspended to 5 mg/µL in 80% methanol.

Chromatographic separation was performed on a UHPLC system equipped with a C18 core-shell column (50 × 2.1 mm, 1.7 µm; Phenomenex). Solvent A was H₂O + 0.1% formic acid, and solvent B was acetonitrile + 0.1% formic acid. The gradient increased from 5% to 50% B over 8 min, then to 99% B by minute 10, followed by washing and re-equilibration. Mass spectrometry was conducted on a Q-Exactive HF using heated electrospray ionization in positive mode. MS1 scans were acquired at 60,000 resolution (120–1500 m/z). Data-dependent MS2 (Top 5) was performed with stepped normalized collision energies of 25/35/45. Data were centroid-transformed with MSconvert and processed with mzmine v4.2.0 [36], and molecular networking was performed in GNPS 2 [37]. Chemical classification was inferred using CANOPUS [38] and SIRIUS [39]. The full feature table is available in Supplementary Table S10 and SIRIUS annotations are available in Supplementary Table S11. Raw data for untargeted metabolomics is available in MassIVE with accession number MSV000096776.

### Biosynthetic gene cluster prediction

Biosynthetic gene clusters (BGCs) in *Corynebacterium propinquum* genomes were predicted using antiSMASH v8.0.2 with default parameters. Predicted clusters were classified according to antiSMASH annotations and similarity to reference clusters in the MIBiG database. Supplementary Table S12 contains the results for the Biosynthetic gene clusters (BGCs) predicted.

### Targeted metabolomics of metabolites

Methanol extracts from the 12-well plates experiment were recovered, centrifuged, and stored at −80 °C until analysis. Metabolites were quantified by targeted LC-MS/MS on an Agilent TQ 6495 using a HILIC-Z column (2.1 × 150 mm, 2.7 µm; Agilent). Data was acquired using a subset of metabolites from the Agilent end-to-end targeted metabolomics workflow as described by Yannell et al. [40]. A table with peak intensities is found in Supplementary Table S13.

### Statistical analyses

Amplicon data were analyzed in *R* v4.4.0 (using RStudio v2024.12.1). SynComs were clustered based on relative abundances at the final time point using partitioning around medoids (PAM) clustering implemented in the *cluster* R package [41]. The optimal number of clusters was automatically determined using the Silhouette method [42], selecting the number of clusters that maximized average silhouette width. Differential metabolite abundance between SynCom clusters was analyzed using the *limma* package [43]. Intensities were log₂-transformed and quantile-normalized. Linear models were fitted for each metabolite, and empirical Bayes moderation was applied to obtain moderated t-statistics, log₂ fold-changes, and Benjamini–Hochberg FDR-adjusted p-values. Metabolites with FDR < 0.05 and |log₂FC| > 1 were considered significantly associated with SynCom clusters. Multi-omics data integration and visualization were performed in R using ggplot2 [44].

## Results

### Nasal SynComs reflect the compositional diversity of the human nasal microbiome

To develop SynComs that resemble the nasal microbiota, we used 5 to 10 bacterial species, each represented by 3 different strains, each isolated from healthy individuals (Figure 1). To identify most prevalent bacteria in the human nasal microbiome, we performed a new analysis of the Human Microbiome Project data from the “nasal cavities” body site (Supplementary Figure S1). We found that the nasal microbiome showed high variability between studied individuals and that the composition is notably uneven, with few species dominating most samples. Additionally, we found that, although species alone did not completely explain *S. aureus* exclusion in the observed samples, samples were grouped in at least broad 4 groups in terms of their composition. Based on this, we selected a pool of 10 bacterial species (*Anaerococcus octavius, Corynebacterium accolens, Corynebacterium propinquum, Corynebacterium pseudodiphtheriticum, Corynebacterium tuberculostearicum, Cutibacterium acnes, Cutibacterium avidum, Dolosigranulum pigrum, Staphylococcus epidermidis, Staphylococcus lugdunensis*) that were prevalent in the human nasal microbiome with the constraint that they were also present in the LaCa collection of nasal bacterial isolates [25].

Next, we assembled 50 SynComs by pooling 5 to 10 randomly selected species and one random strain per species and growing them in synthetic nasal medium (Figure 1). To evaluate the stability and assembly process of the SynComs, they were cultured iteratively 4 times with reinoculations in fresh plates every 3 days to ensure sufficient supply with nutrients. In the last reinoculation, we added *S. aureus* to test the colonization resistance of the SynComs. We characterized the composition of the communities at the final time point (72 h after *S. aureus* was added, Figure 1). Across randomly assembled communities, we observed variable end-point composition, with some SynComs converging toward stable multi-species composition while others became dominated by a single species (Figure 2). The SynComs were grouped in three clusters (compositional similarity with silhouette method) with different species richness, evenness, dominant species and different resistance to *S. aureus* colonization (Figure 2). Importantly, SynComs with similar inocula at the species-level resulted in different end-point compositions, thus indicating a large influence of strain-level variation. For instance, SynComs that contained all the 10 species (Supplementary Table S2) with different strains that lead to different final compositions and cluster belonging included SC1 (Cluster 1), SC29 (Cluster 2), SC5 (Cluster 3).

**Figure 2.**
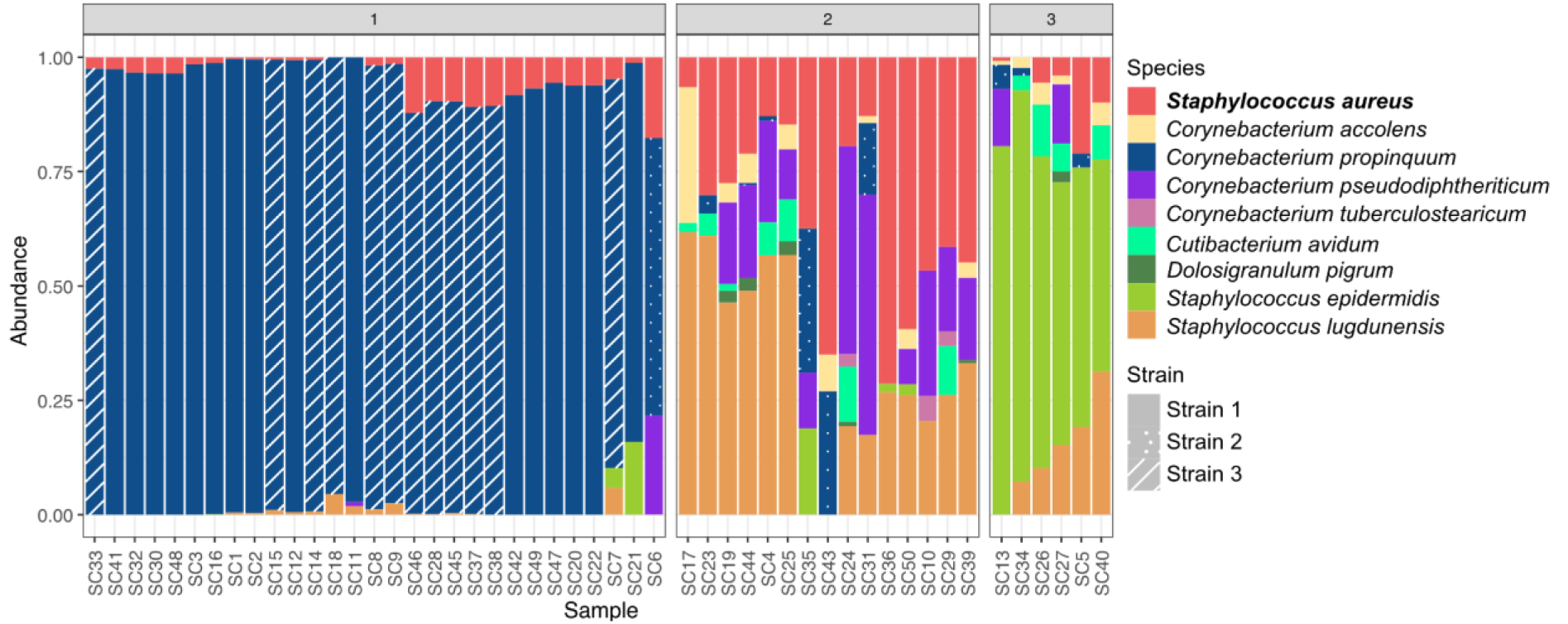
Screening of SynCom composition and assembly into stable clusters. Relative abundance barplots show the microbial composition of the 50 assembled SynComs at the final study time point (72 h after *S. aureus* inoculation). To screen the ecological behavior across the broad compositional landscape of the SynComs, one replicate per SynCom was characterized. Samples are ordered along the x-axis according to Euclidean-distance clustering of relative abundance data. The optimal number of clusters (k=3) was determined using the silhouette method, identifying distinct compositional attractors. Strain-level variation is specifically highlighted for *C. propinquum* and *D. pigrum* using distinct barplot patterns.

The end-point composition of the 50 SynComs clustered in three distinct groups. Cluster 1 was dominated by *C. propinquum* (mean of 93.1%) and showed high resistance to *S. aureus* (mean abundance of 4.6%, Figure 2). Just a few other species occurred at low abundance in cluster 1 SynComs (*e.g.*, *Staphylococcus lugdunensis* and *S. epidermidis*). Thus, the 20 SynComs of cluster 1 consisted primarily in co-cultures of *C. propinquum* and *S. aureus*. The other two clusters showed higher species diversity. Cluster 2 frequently included *S. lugdunensis*, *C. pseudodiphtheriticum*, *C. avidum*, *D. pigrum*, *C. accolens*, and *C. tuberculostearicum*. This cluster had the lowest colonization resistance, with a mean relative abundance of *S. aureus* of 34.1%. Cluster 3 included similar species as cluster 2 but was dominated by *S. epidermidis* and *S. lugdunensis*. Cluster 3 showed strong colonization resistance, with a mean final relative abundance of *S. aureus* of 6.9%. Two bacterial species *A. octavius* and *C. acnes* were not detected in any SynCom.

### Strain-level diversity influenced population dynamics and resistance to S. aureus colonization

Next, we selected 20 representative SynComs that captured the compositional diversity of the full data set. We characterized their composition at all time points, which included the 4 samples taken every 72 h when sampling for biomass to reinoculate the SynComs in fresh SNM3 plates. For these SynComs, we sequenced all time points and replicates. Figure 3 shows one replicate per SynCom and time point. Supplementary Figure S2.A shows the Bray-Curtis distance between the replicates which was below 0.2 for most of the SynComs, showing reproducibility of our experimental setup.

**Figure 3.**
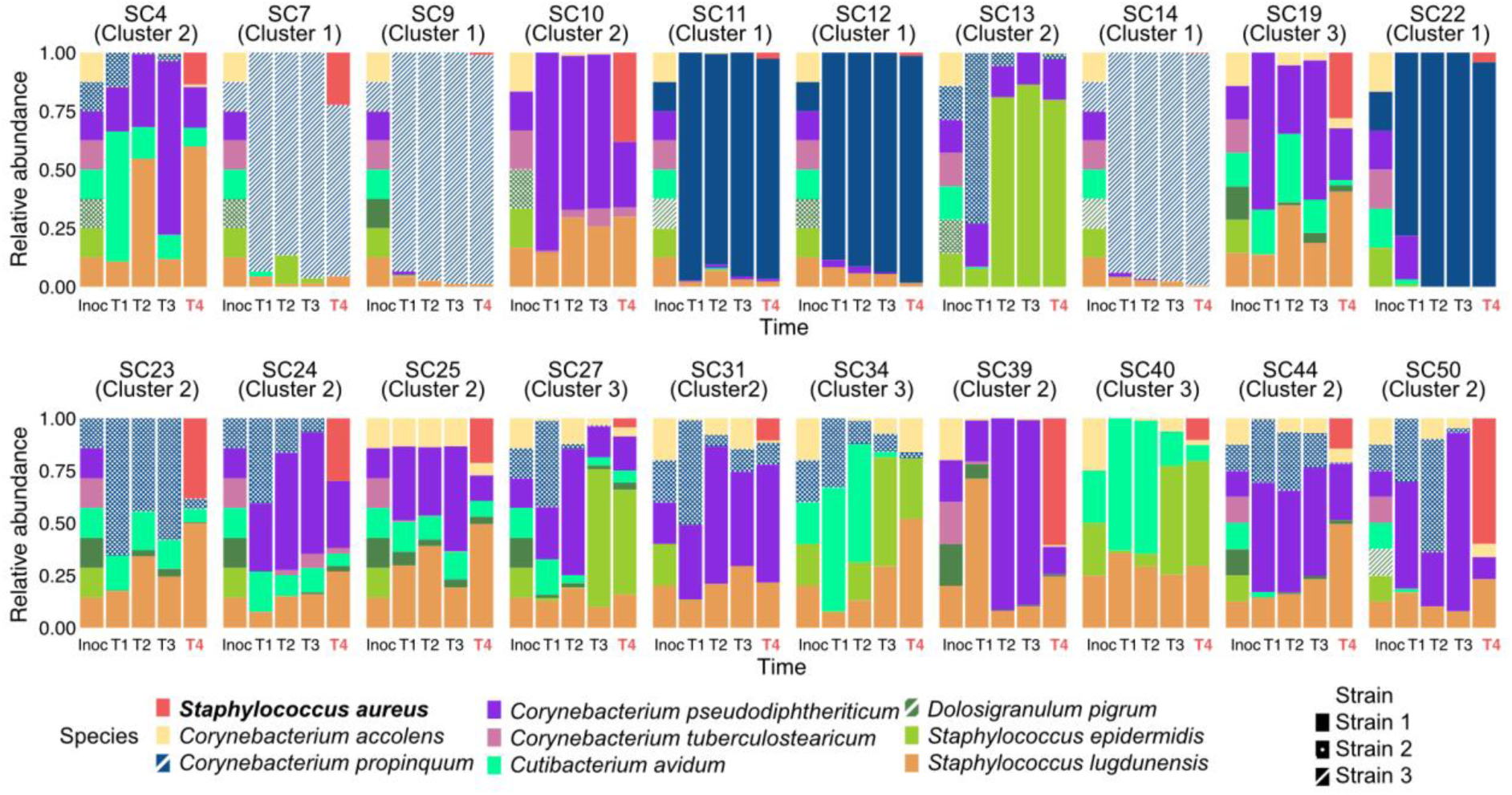
Temporal dynamics of bacterial community composition in 20 selected SynComs. Samples of 20 SynComs were selected out of the 50 in the main experiment and community composition was characterized by sequencing at four time points across the experiment. Samples from these 20 SynComs were also analyzed using untargeted metabolomics. Each panel represents one SynCom and shows relative abundance barplots for one replicate per SynCom over time. Barplot patterns indicate which of the three strains was inoculated for *C. propinquum* and *D. pigrum*. Strain level information of other species is not shown.

Sequencing of the 20 SynComs revealed that both species and strain identity had a substantial impact on SynCom assembly and composition. In addition to Figure 3, Supplementary Figure S2.B shows the Bray-Curtis distance between the first three timepoints and the final composition for all sequenced SynComs. In general, there were two types of assembly patterns, one that stabilized quickly after timepoint 1 and another that varied at the 4 time points. The first group (SynComs 7, 9, 11, 12, 14 and 22) with fast stabilization included SynComs that were dominated by *C. propinquum* strains 16 and 265. These SynComs showed low dissimilarity values from the beginning of the experiment compared to the final state. The other SynComs showed less stable dynamics with a more fluctuating composition and higher dissimilarity values in respect to the final state, which was dependent on the species and strains inoculated.

Next, we observed that strain-level variation played an important role for assembly of the different SynComs. For example, the presence of distinct strains of the same species had a strong effect on the abundance of *S. aureus* at the final time point. The strongest strain-level effect observed was the case of *C. propinquum* in cluster 1, in which all except for one of the SynComs consisted of two *C. propinquum* strains (Nr. 16 and Nr. 265; Figure 2). In contrast, *C. propinquum* strain 70 was present only during the initial time points and other species overgrew it by the end of the experiment (for example SynComs 13, 23 and 50, Figure 3). Thus, *C. propinquum* strain 70 only dominated in SynCom 6 (Figure 2). Similarly, *D. pigrum* strain 21 was the only strain of this species that could persist through the entire cultivation period in the SynComs where it was included, although its abundance was always below 5%. *D. pigrum*, *A. octavius* and *C. acnes* are fastidious bacteria and neither of their strains showed growth when we tried growing them as monocultures in SNM3 (Supplementary Figure S3). Thus, in the case of *D. pigrum*, strain 21 was able to grow only when it was part of these specific SynComs when growing in SNM3, suggesting that it obtains essential nutrients or cofactors from other bacteria in these communities. Additionally, we observed co-occurrence patterns at species level such between *C. accolens*, *C. avidum*, *C. pseudodiphtheriticum*, and *D. pigrum*, independently of the strains used for the first two species (for example in SynComs 19, 25, 27, 44 and 50).

We then performed co-cultures between each of the 3 strains of *C. propinquum* and *S. aureus* USA300 to explore how the nutritional environment can affect the interaction between these two species. *C. propinquum* strains 16 and 265 (but not strain 70) were able to dominate over *S. aureus* under the nutritional conditions of SNM3 (Supplementary Figure S4). In BHI medium, in contrast, *S. aureus* dominated the pairwise co-cultures against all three *C. propinquum* strains (Supplementary Figure S4). Additionally, the *C. propinquum* strains 16 and 265 achieved higher optical densities after 24 h growth in liquid medium than strain 70 (Supplementary Figure S3).

### Metabolome of SynComs differs between clusters and suggests nutritional competition

To investigate the metabolism of the different SynComs, we performed untargeted metabolomics on all time points and replicates of the 20 selected SynComs (Figure 4). Metabolite extracts in methanol:water were collected from plates and analyzed by untargeted LC–MS/MS enabling comprehensive detection of small-molecule features without prior assumptions about metabolite identities.

**Figure 4.**
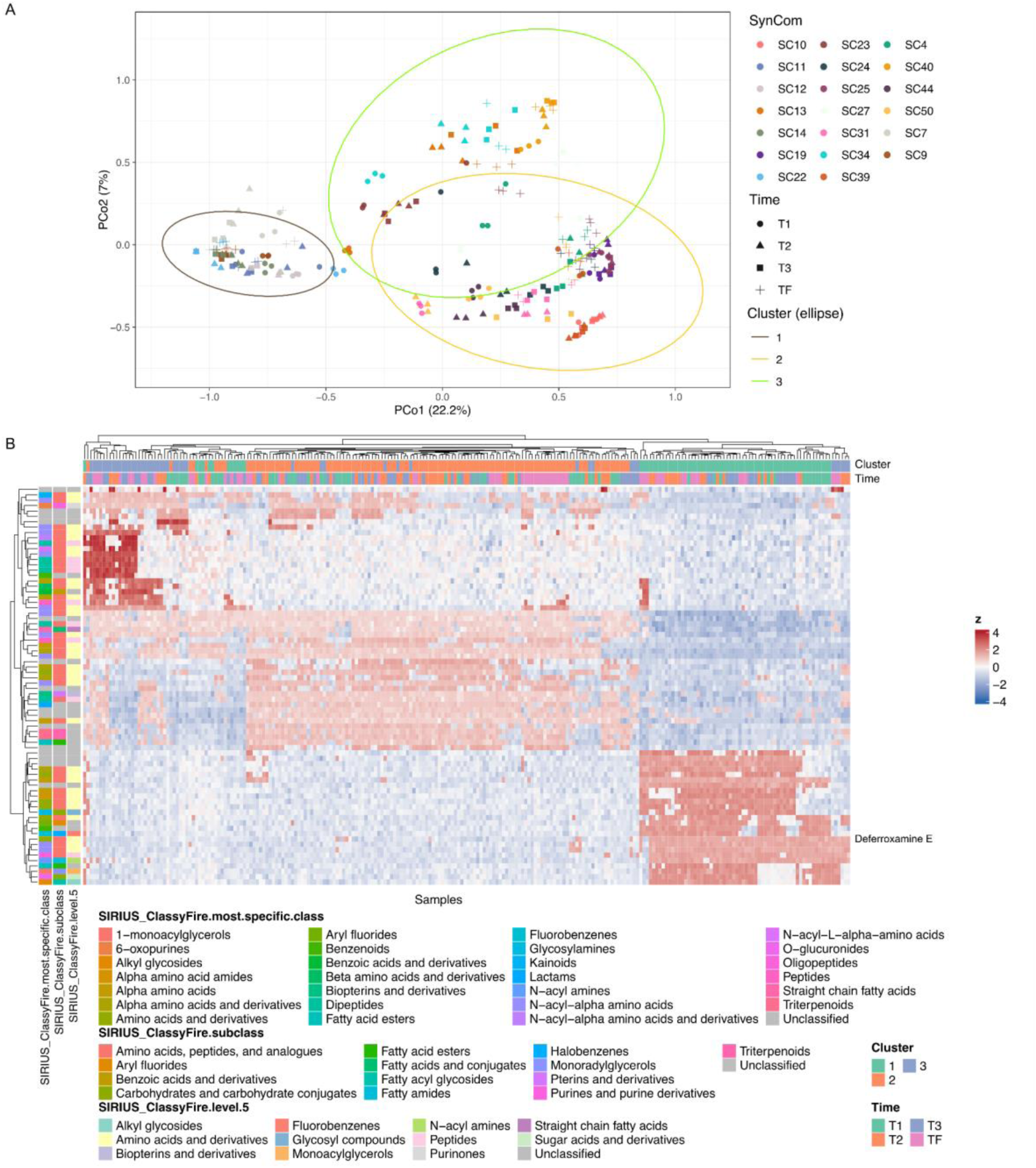
Compositional similarity and metabolic profiles of the 20 selected SynComs. For 20 of the original 50 SynComs, community composition was characterized and untargeted metabolomics analyses were performed at four time points across the experiment and 3 replicates (240 samples). (A) Principal coordinates analysis (PCoA) of bacterial community composition for samples from the 20 selected SynComs. The ordination is based on pairwise dissimilarities in relative abundances. Each point represents one sample and replicate (3 per SynCom in each timepoint); colors indicate SynCom identity, point shapes indicate the time point from which the samples were taken and 95% confidence ellipses denote the SynCom composition clusters. (B) Heatmap of the top 30 metabolite features associated with SynCom clusters, identified using a limma-based linear modeling approach using the limma package for R [40]. The column annotation above the heatmap indicates the cluster assignment of each sample and the time point from which they were taken. Row annotations on the left indicate predicted metabolite classes as assigned by the SIRIUS CANOPUS algorithm. Metabolite signals are displayed as standardized values (row-wise z-scores of log-transformed intensities).

We used Linear Models for Microarray Data (limma) to test whether the identified SynCom clusters according to microbial composition (Figure 2 and Figure 4A) differed in their metabolic profiles. Limma applies linear modeling and an empirical Bayes method to detect features that differ significantly between predefined groups while accounting for variance and multiple testing. This analysis showed that each cluster of SynComs has a distinct metabolic signature (Figure 4B). Especially SynComs from Cluster 1, which were dominated by *C. propinquum*, showed enrichment of specific metabolites such as the siderophore deferoxamine E, indicating that this metabolite is associated with high abundance of *C. propinquum* in these communities. In the genomes of the three *C. propinquum* strains, seven biosynthetic gene clusters (BGCs) were predicted using antiSMASH. In particular, one high-confidence NI-siderophore cluster showed strong similarity to the dehydroxynocardamine biosynthetic pathway, consistent with the detection of deferoxamine E in the untargeted metabolomics data. Annotation with SIRIUS predicted that the other compounds associated with the SynCom clusters are mainly amino acids, their derivatives, and peptides.

### Metabolites production and consumption profiles are SynCom-specific and influence S. aureus colonization resistance

To assess the role of amino acids and other metabolites on the nutritional competition in community assembly and *S. aureus* colonization resistance, we analyzed a panel of 24 nutrients involved in primary metabolism of bacteria, including amino acids, nucleosides, and vitamins for 5 selected SynComs (Figure 5 and Supplementary Table S10). These 5 SynComs were representative from clusters 1 (SC7 and SC12) and cluster 3 (SC27 and SC40) that showed high *S. aureus* colonization resistance and 1 SynCom from cluster 2 (with low *S. aureus* colonization resistance). Importantly, when we cultured these 5 SynComs for this metabolomics experiment, their composition matched the results from the main experiment, demonstrating the reproducibility of the cultivation protocol and SynCom stability (Figure 5A). Additionally, we measured the 24 compounds in monocultures of *S. aureus* USA300 and the three *C. propinquum* strains used in the SynCom experiments. Using 12-well agar plates, we quantified concentration changes of the 24 nutrients relative to their starting concentrations in the medium (measured in control wells without inoculum; Figure 5B and 5C). These data showed that some nutrients are increasing relative to the medium, indicating that they are produced by specific bacteria. For example, all 5 SynComs and the three *C. propinquum* strains had high levels of riboflavin and tyrosine, which were hardly measurable in the medium and in *S. aureus* cultures.

**Figure 5.**
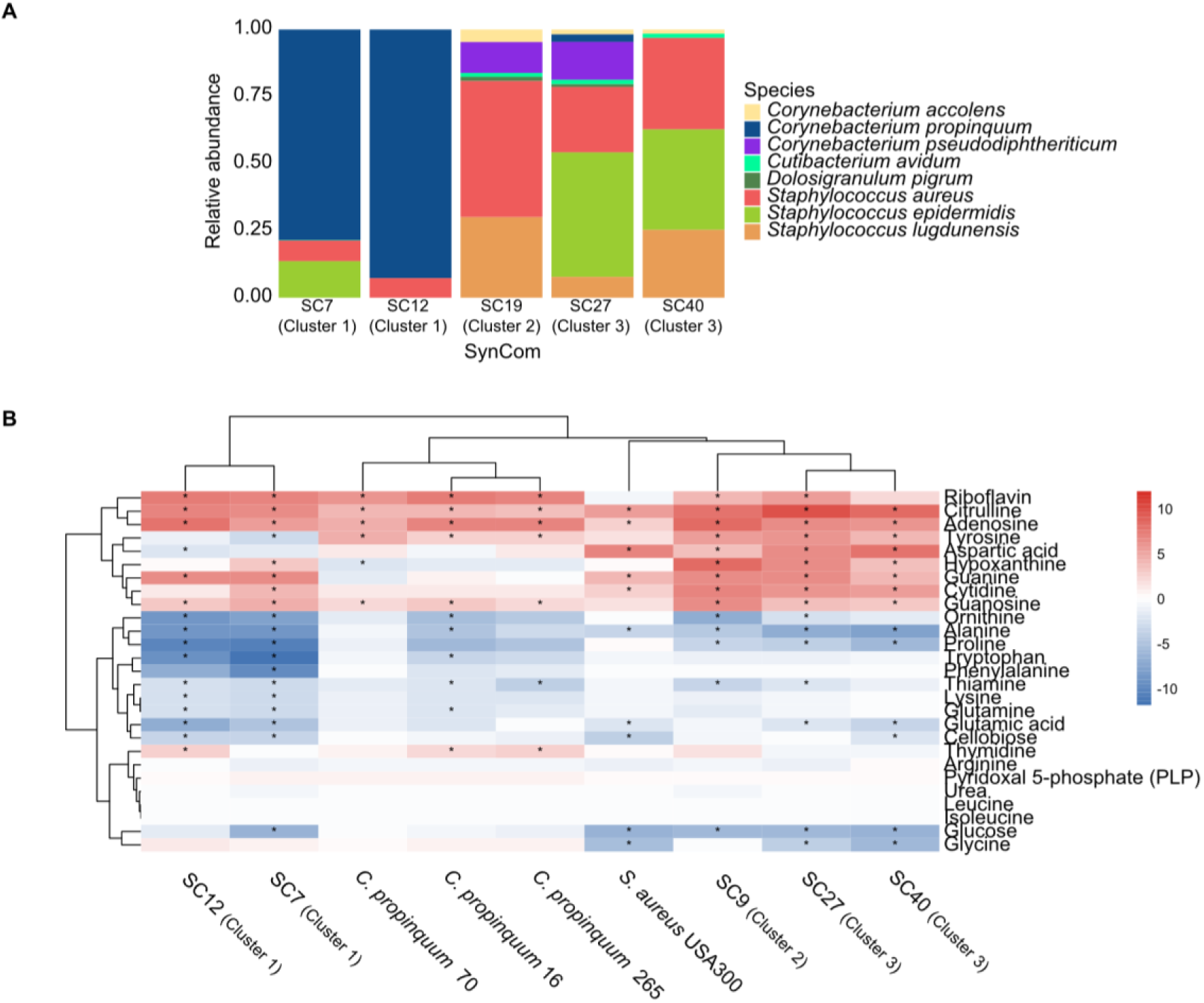
Bacterial composition and targeted metabolite profiles of five selected SynComs, *S. aureus* USA300 and the three strains of *C. propinquum* used in SynCom experiments. Five SynComs selected from the main screen were grown again in 12-well plates to quantify metabolite production and consumption by targeted metabolomics. In parallel, monocultures of *S. aureus* USA300 (used to test colonization resistance) and the three *C. propinquum* strains were cultivated in the same format and analyzed alongside the SynComs. (A) Relative abundance barplots showing the bacterial composition of the five selected SynComs. (B) Heatmap of log₂ fold-change in mean metabolite abundance relative to the medium-only control (SNM3 without inoculation). For each metabolite and condition, log₂ fold-changes were computed from mean intensities across replicates. Asterisks indicate metabolites that differ significantly from the control based on Welch’s t-tests on log₂-transformed replicate intensities with Benjamini–Hochberg correction (FDR < 0.05) and an absolute log₂ fold-change ≥ 2 (≥ 4-fold change).

Other nutrients were depleted in *S. aureus* cultures relative to the medium, such as alanine, glutamic acid, and glycine. SynComs 10 and 12, which were dominated by *C. propinquum*, showed strong decreases of several nutrients (at least 4-fold change in 10 out of 24 metabolites measured) including ornithine, Alanine, Proline, Tryptophane, Phenylalanine, thiamine, Lysine, glutamine, glutamic acid (Figure 5B). Similarly, SynComs 20, 27, and 40 showed high consumption of amino acids such as glycine, alanine, and glutamic acid.

## Discussion

In this study, we assembled synthetic communities (SynComs) from 10 prevalent human nasal bacterial species (*Anaerococcus octavius, Corynebacterium accolens, Corynebacterium propinquum, Corynebacterium pseudodiphtheriticum, Corynebacterium tuberculostearicum, Cutibacterium acnes, Cutibacterium avidum, Dolosigranulum pigrum, Staphylococcus epidermidis, and Staphylococcus lugdunensis*). Our unbiased assembly strategy allowed us to test whether nasal community state types arise as intrinsic ecological attractors. Additionally, we evaluated how community composition and strain-level variation influence stability of these SynComs over time (four time points across 12 days) and resistance to invasion of *S. aureus* at the final time point.

Multistability has been documented across microbial communities, where interaction-driven population dynamics and feedback loops generate alternative taxonomic states [45, 46]. In the case of the nasal microbiome, community state types (CSTs) have been described based on a species level [26]. However, comparison with the HMP dataset suggests that CST boundaries are diffuse, indicating that additional sources of variability shape nasal community composition than previously thought. In the case of our SynComs experiment, three distinct community clusters emerged despite random assembly, suggesting that deterministic ecological processes play an important role in community dynamics. These clusters can be interpreted as three alternative stable states, each representing similar community compositions towards which the different combinations of species and strains tend to converge.

The clusters differed in species richness, evenness, and dominant taxa, and showed markedly different levels of resistance to *S. aureus* colonization. Importantly, our panel of cultivated SynComs reproduced several characteristics observed in the HMP dataset and the previously described CSTs of the nasal microbiome (Figure S1). In particular, we observed the frequent coexistence of *Corynebacterium* species, the persistence of *D. pigrum* (although represented by a single strain), and the overall unevenness typical of nasal communities. Together, these findings demonstrate that the ability of the nasal microbiome to resist *S. aureus* colonization can be recapitulated and studied under controlled *in vitro* conditions, making our SynComs a platform for systematic and mechanistic analyses of the nasal microbiota dynamics.

The observation that SynComs dominated by *C. propinquum* showed stronger colonization resistance against *S. aureus* is consistent with previous studies showing that nasal communities dominated by *Corynebacterium* spp. (including *C. propinquum*) are often associated with reduced *S. aureus* carriage, potentially through production of siderophores and other competitive factors that limit *S. aureus* growth [8, 11, 13, 47]. Analysis of the HMP data supports this hypothesis and shows that species identities are not the only predictive factor of colonization resistance. Our SynCom experiments show three examples of strain-level effects: (1) SynComs with the same species composition, but with different strains, resulted in different final compositions (for example SC1 of Cluster 1, SC29 of Cluster 2, SC5 of Cluster 3); (2) *C. propinquum* strains 16 and 265 consistently dominated SynComs, whereas strain 70 did not and (3) *D. pigrum* strain 21 was the only strain of that species which could persist throughout the entire experiment. Together with the high variability of *C. propinquum* abundance across the HMP cohort, ranging from very low relative abundance to >70% in some individuals, these findings suggest that strain-level characteristics are important for nasal microbiome composition and function. Our data shows that differences between strains (presumably their metabolic capacities) influences their ability to dominate and persist. Thus, future studies could map this to specific metabolic capacities of the different strains, e.g. biosynthetic pathways for amino acids or siderophores. For example, it has been shown that mutations in biosynthetic genes can lead to strain-specific auxotrophies [48]. The same is true for other species, such as *S. lugdunensis* or *D. pigrum*, for which only some strains have shown inhibitory capacities against *S. aureus* [9, 49, 50].

*D. pigrum* is another key species in the nasal environment, ranking as the eighth most prevalent species in the HMP cohort data. It has been noted as promoting colonization resistance through modulation of the local microenvironment and production of antimicrobial factors [9, 50]. Moreover, *D. pigrum* has been reported to engage in positive interactions with *Corynebacterium* species, particularly *C. pseudodiphtheriticum* [9, 51]. Our SynCom data support these observations, as *D. pigrum* was only present in SynComs when *C. pseudodiphtheriticum* and *C. accolens* were also present (SynComs 19, 25, 27, 44 and 50; Figure 2). While the mechanistic basis of these associations remains unclear, understanding such cooperative interactions is important given their potential synergistic effects against pathogens such as *S. aureus*.

In addition to Corynebacteria and *D. pigrum*, staphylococcal species, such as *S. epidermidis* and *S. lugdunensis*, are known to inhibit *S. aureus* through production of antimicrobial peptides and direct interference [49, 52–54]. In our SynComs, rather than observing isolated effects of individual species, we found evidence for synergistic interactions between these two staphylococci due to their consistent coexistence in the SynComs (SynComs 5, 7, 21, 26, 27, 34, 36, 40, 50; Figure 2). Such potential cooperative effects have been described in previous research in the context of the nasal environment and the respiratory tract, for example, synergism between *D. pigrum* and *C. pseudodiphtheriticum* [51], or siderophore sharing between staphylococcal species [47]. These results suggest that colonization resistance in the nasal cavity is likely an emergent property of multiple interacting species and strains, and probably driven by both competition and cooperation.

Our study also shows that it is possible to quantify both primary and secondary metabolites in nasal SynComs. These metabolome analyses provide additional insights into potential mechanisms underlying colonization resistance. SynComs in Cluster 1, dominated by *C. propinquum*, were enriched in the siderophore deferoxamine E. Genome analysis further revealed a high-confidence NI-siderophore biosynthetic gene cluster with strong similarity to the dehydroxynocardamine pathway, consistent with the production of deferoxamine E. This siderophore may contribute to iron depletion and thereby limit *S. aureus* growth. Previous studies have demonstrated that iron sequestration by siderophores is an important competitive mechanism of corynebacteria in nasal ecosystems [13, 47, 55, 56]. Moreover, we observed changes in the abundance of nutrients, such as amino acids and riboflavin, that were highly specific for some SynComs, suggesting that nutrient competition may further contribute to colonization resistance. We identified alanine, glutamic acid, tyrosine, glycine and riboflavin as important nutritional metabolites for *S. aureus* USA300. These results are consistent with studies reporting that *S. aureus* growth is constrained by amino acid availability in nutrient-limited environments and by the fact that the nasal cavity is naturally poor in amino acids [10, 25, 55].

Our pairwise coculture experiments with *C. propinquum* and *S. aureus* further support the hypothesis that nutritional conditions influence the competitive dynamics between the two bacteria. Under the poor nutritional conditions of SNM3, *C. propinquum* strains 16 and 265 successfully outcompeted *S. aureus*. In contrast, the nutrient-rich medium BHI allowed *S. aureus* to overgrow all *C. propinquum* strains. These contrasting outcomes suggest that differences in nutrient availability shape the relative fitness of the two species.

One possible explanation is that *C. propinquum* is adapted to the nutritionally limited environment of the nasal cavity, potentially favoring efficient resource utilization, while *S. aureus* may be better suited to rapid growth under nutrient-rich conditions. Although this hypothesis remains to be directly tested, it is consistent with expectations from the r/K selection theory [57] where the slow-growing K-strategists (in this case *C. propinquum* strains) specialize in surviving under poor-nutrient conditions and r- strategists (*S. aureus* in this scenario) maximize their growth rate in nutrient rich scenarios. r/K strategies have been observed in other microbial systems, like soil communities, as important for community structure and functioning [58, 59]. Further, this context-dependent competitive outcome is a widely reported phenomenon, demonstrating that competition for specific limiting nutrients, such as essential metal cofactors or amino acids, mediates colonization resistance. An example of this was reported as a colonization resistance mechanism in the gut microbiome against *Salmonella* [60]. Additionally, our observation that strains 16 and 265 grow better than strain 70 both in single and in SynCom cultures suggests that intrinsic strain-level differences are important factors of competitive success. Together, these results indicate that both environmental constraints (e.g. nutrient availability) and strain specific factors (e.g. biosynthetic or catabolic capacities) determine the ability of *C. propinquum* to inhibit *S. aureus*, providing an explanation for the strain-dependent protection observed in our SynCom experiments.

The reproducibility of our SynCom approach shows that they are a tractable model system for controlled and systematic investigations of nasal microbiome ecology. Until now, most studies examining the effects of specific taxa on *S. aureus* have used isolated pairs of strains, and here we expand this to a broader community context. Our work highlights how strain-level diversity, species co-occurrence, and metabolic interactions collectively shape colonization outcomes.

### Limitations and Future Directions

Our study has several limitations. First, our SynComs were assembled from a limited culture collection of 10 species, which, while representing some of the most prevalent members of the nasal microbiome, does not capture its full diversity. Fastidious or currently unculturable taxa, not present in the LaCa collection, were necessarily excluded. Our experimental setting did not reproduce the anaerobic microenvironments present in the nasal cavity, and as a result two (*A. octavius* and *C. acnes*) of the three anaerobic species failed to grow in any SynCom. These anaerobes have been implicated in community structure and may influence *S. aureus* dynamics [61]. Furthermore, only one strain of *S. aureus* was used in our invasion test. Use of other strains of *S. aureus* could help elucidate the strain-dependent mechanisms of invasion success for this species.

Second, our experiments were performed entirely *in vitro*, which does not fully recapitulate the nasal mucosal environment, host immune responses, or spatial structure that may modulate microbial interactions and metabolite diffusion, as has been shown for other microbial ecosystems. Future studies should expand SynCom designs to include additional, fastidious species and incorporate host-derived factors such as epithelial cells, immune components, or human organoids, as has been done in porcine nasal microbiome models [62]. Incorporating physical structure into experimental setups is necessary, as it could help establish microaerophilic or anaerobic niches that support the growth of strict anaerobes and more closely mimic in vivo spatial organization which influences ecological interactions and dynamics [63, 64]. Finally, while our metabolomics analysis identified metabolites associated with specific community clusters, a substantial proportion of detected features remain unannotated, which limits mechanistic interpretation. Future work combining targeted metabolomics and genetic perturbations will be essential to confirm the specific roles of siderophores and amino acid competition in mediating *S. aureus* exclusion.

In summary, we demonstrate that nasal SynComs reproduce key compositional and functional features of the human nasal microbiome, including the emergence of distinct community states with variable resistance to *S. aureus* colonization. Our findings highlight the importance of both species and strain identity in shaping community assembly and function and suggest that nutritional competition particularly for iron and amino acids is a critical mechanism driving colonization resistance in a community context. These results provide a basis for the rational design of microbiome-based interventions aimed at reducing *S. aureus* carriage and preventing infection. By integrating SynCom approaches with host-microbe models and *in vivo* validation, future research can further elucidate the ecological principles governing nasal microbiome stability and pathogen exclusion.

## Acknowledgements

We thank Dr. Anna Walke from Agilent Technologies (Germany, Waldbronn) for her guidance on the targeted LC-MS/MS Agilent method. This work was funded by the Deutsche Forschungsgemeinschaft (DFG, German Research Foundation) under Germanýs Excellence Strategy – EXC 2124 – 390838134 (M.N.D, S.H., D.P., L.C., L.T.A., H.L., S.He.).

## Data availability

Data and code for analyses are available on GitHub at: https://github.com/marcelo-nd/nasalSynComs.

## References

1. Caballero-Flores G, Pickard JM, Núñez G. Microbiota-mediated colonization resistance: mechanisms and regulation. Nat Rev Microbiol 2023;21:347–360. 10.1038/s41579-022-00833-7

2. Sarkar S et al. A Review on the Nasal Microbiome and Various Disease Conditions for Newer Approaches to Treatments. Indian J Otolaryngol Head Neck Surg 2023;75:755–763. 10.1007/s12070-022-03205-y

3. Sakr A et al. Staphylococcus aureus Nasal Colonization: An Update on Mechanisms, Epidemiology, Risk Factors, and Subsequent Infections. Front Microbiol 2018;9:2419. 10.3389/fmicb.2018.02419

4. Eiff CV, Peters G. Nasal Carriage as a Source of Staphylococcus aureus Bacteremia. 2001.

5. Duhaniuc A et al. Multidrug-Resistant Bacteria in Immunocompromised Patients. Pharmaceuticals 2024;17:1151. 10.3390/ph17091151

6. Chand U, Priyambada P, Kushawaha PK. Staphylococcus aureus vaccine strategy: Promise and challenges. Microbiol Res 2023;271:127362. 10.1016/j.micres.2023.127362

7. Van Dalen R et al. Secretory IgA impacts the microbiota density in the human nose. Microbiome 2023;11:233. 10.1186/s40168-023-01675-y

8. Laux C, Peschel A, Krismer B. *Staphylococcus aureus* Colonization of the Human Nose and Interaction with Other Microbiome Members. Microbiol Spectr 2019;**7**:7.2.34. 10.1128/microbiolspec.GPP3-0029-2018

9. Brugger SD et al. Dolosigranulum pigrum Cooperation and Competition in Human Nasal Microbiota. mSphere 2020;5:e00852–20. 10.1128/mSphere.00852-20

10. Krismer B et al. Nutrient Limitation Governs Staphylococcus aureus Metabolism and Niche Adaptation in the Human Nose. PLoS Pathog 2014;10:e1003862. 10.1371/journal.ppat.1003862

11. Tamkin E et al. Airway *Corynebacterium* interfere with *Streptococcus pneumoniae* and *Staphylococcus aureus* infection and express secreted factors selectively targeting each pathogen. Infect Immun 2025;93:e00445–24. 10.1128/iai.00445-24

12. Krismer B et al. The commensal lifestyle of Staphylococcus aureus and its interactions with the nasal microbiota. Nat Rev Microbiol 2017;15:675–687. 10.1038/nrmicro.2017.104

13. Zhao Y et al. Nasal commensals reduce *Staphylococcus aureus* proliferation by restricting siderophore availability. ISME J 2024;18:wrae123. 10.1093/ismejo/wrae123

14. Proctor DM, Relman DA. The Landscape Ecology and Microbiota of the Human Nose, Mouth, and Throat. Cell Host Microbe 2017;21:421–432. 10.1016/j.chom.2017.03.011

15. Hardy BL, Merrell DS. Friend or Foe: Interbacterial Competition in the Nasal Cavity. J Bacteriol 2021;203. 10.1128/JB.00480-20

16. Faith JJ et al. The Long-Term Stability of the Human Gut Microbiota. Science 2013;341:1237439. 10.1126/science.1237439

17. Venturelli OS et al. Deciphering microbial interactions in synthetic human gut microbiome communities. Mol Syst Biol 2018;14:e8157. 10.15252/msb.20178157

18. Bai Y et al. Functional overlap of the Arabidopsis leaf and root microbiota. Nature 2015;528:364–369. 10.1038/nature16192

19. Niu B et al. Simplified and representative bacterial community of maize roots. Proc Natl Acad Sci 2017;114. 10.1073/pnas.1616148114

20. Goodman AL et al. Extensive personal human gut microbiota culture collections characterized and manipulated in gnotobiotic mice. Proc Natl Acad Sci 2011;108:6252–6257. 10.1073/pnas.1102938108

21. Pacheco AR, Osborne ML, Segrè D. Non-additive microbial community responses to environmental complexity. Nat Commun 2021;12:2365. 10.1038/s41467-021-22426-3

22. Weiss AS et al. In vitro interaction network of a synthetic gut bacterial community. ISME J 2022;16:1095–1109. 10.1038/s41396-021-01153-z

23. Buffie CG et al. Precision microbiome reconstitution restores bile acid mediated resistance to Clostridium difficile. Nature 2015;517:205–208. 10.1038/nature13828

24. Bonillo-Lopez L et al. *In vitro* metabolic interaction network of a rationally designed nasal microbiota community. 2024. Systems Biology, 2024.

25. Camus L et al. Tyrosine availability shapes Staphylococcus aureus nasal colonization and interactions with commensal communities. bioRxiv 2025;2025.05.06.651429. 10.1101/2025.05.06.651429

26. Liu CM et al. *Staphylococcus aureus* and the ecology of the nasal microbiome. Sci Adv 2015;1:e1400216. 10.1126/sciadv.1400216

27. Turnbaugh PJ et al. The Human Microbiome Project. Nature 2007;449:804–810. 10.1038/nature06244

28. Estaki M et al. QIIME 2 Enables Comprehensive End-to-End Analysis of Diverse Microbiome Data and Comparative Studies with Publicly Available Data. Curr Protoc Bioinforma 2020;70:e100. 10.1002/cpbi.100

29. Callahan BJ et al. DADA2: High-resolution sample inference from Illumina amplicon data. Nat Methods 2016;13:581–583. 10.1038/nmeth.3869

30. Camacho C, et al. BLAST+: architecture and applications. BMC Bioinformatics 2009;10:421. 10.1186/1471-2105-10-421

31. Goldfarb T et al. NCBI RefSeq: reference sequence standards through 25 years of curation and annotation. Nucleic Acids Res 2025;53:D243–D257. 10.1093/nar/gkae1038

32. Aguayo-Reyes A et al. Emergence of ST8-USA300 and ST8-USA300-Latin American variant: a changing landscape of community-associated methicillin-resistant *Staphylococcus aureus* in Chile. Microbiol Spectr 2025;13:e01031–25. 10.1128/spectrum.01031-25

33. Seol D et al. Microbial Identification Using rRNA Operon Region: Database and Tool for Metataxonomics with Long-Read Sequence. Microbiol Spectr 2022;10:e02017–21. 10.1128/spectrum.02017-21

34. Curry KD et al. Emu: species-level microbial community profiling of full-length 16S rRNA Oxford Nanopore sequencing data. Nat Methods 2022;19:845–853. 10.1038/s41592-022-01520-4

35. Rice P, Longden I, Bleasby A. EMBOSS: The European Molecular Biology Open Software Suite. Trends Genet 2000;16:276–277. 10.1016/S0168-9525(00)02024-2

36. Schmid R et al. Integrative analysis of multimodal mass spectrometry data in MZmine 3. Nat Biotechnol 2023;41:447–449. 10.1038/s41587-023-01690-2

37. Nothias L-F et al. Feature-based molecular networking in the GNPS analysis environment. Nat Methods 2020;17:905–908. 10.1038/s41592-020-0933-6

38. Dührkop K et al. Systematic classification of unknown metabolites using high-resolution fragmentation mass spectra. Nat Biotechnol 2021;39:462–471. 10.1038/s41587-020-0740-8

39. Dührkop K et al. SIRIUS 4: a rapid tool for turning tandem mass spectra into metabolite structure information. Nat Methods 2019;16:299–302. 10.1038/s41592-019-0344-8

40. Yannell, Karen, Hsiao, Jordy, Cuthbertson, Dan. Mastering HILIC-Z Separation for Polar Analytes. 2023. USA: Agilent Technologies, Inc., 2023.

41. Maechler M, et al. cluster: Cluster analysis basics and extensions (R package version 2.1.1). 2021. 2021.

42. Rousseeuw PJ. Silhouettes: A graphical aid to the interpretation and validation of cluster analysis. J Comput Appl Math 1987;20:53–65. 10.1016/0377-0427(87)90125-7

43. Ritchie ME et al. limma powers differential expression analyses for RNA-sequencing and microarray studies. Nucleic Acids Res 2015;43:e47–e47. 10.1093/nar/gkv007

44. Wickham H. ggplot2 Elegant Graphics for Data Analysis, 2nd edn. Springer Cham, 2016.

45. Gonze D et al. Multi-stability and the origin of microbial community types. ISME J 2017;11:2159–2166. 10.1038/ismej.2017.60

46. Estrela S et al. Functional attractors in microbial community assembly. Cell Syst 2022;13:29–42.e7. 10.1016/j.cels.2021.09.011

47. Stubbendieck RM et al. Competition among Nasal Bacteria Suggests a Role for Siderophore-Mediated Interactions in Shaping the Human Nasal Microbiota. Appl Environ Microbiol 2019;85:e02406–18. 10.1128/AEM.02406-18

48. Lubrano P et al. Metabolic mutations reduce antibiotic susceptibility of E. coli by pathway-specific bottlenecks. Mol Syst Biol 2025;21:274–293. 10.1038/s44320-024-00084-z

49. Heilbronner S. Staphylococcus lugdunensis. Trends Microbiol 2021;29:1143–1145. 10.1016/j.tim.2021.07.008

50. Cole AL et al. Nasal microbiome inhabitants with anti- *Staphylococcus aureus* activity. Microbiol Spectr 2025;e01024–25. 10.1128/spectrum.01024-25

51. Cisneros M et al. Synergistic inhibition of pneumococcal growth by *Dolosigranulum pigrum* and *Corynebacterium pseudodiphtheriticum:* insights into nasopharyngeal microbial interactions. Microbiol Spectr 2025;13:e00138–25. 10.1128/spectrum.00138-25

52. Liu C-C et al. Defects in energy metabolism increase the susceptibility of *Staphylococcus aureus* and its small colony variants (SCVs) to *Staphylococcus lugdunensis* and lugdunin. Microbiol Spectr 2025;e01006–25. 10.1128/spectrum.01006-25

53. Bessesen MT. Interventions targeting the nasal microbiome to eradicate methicilin-resistant Staphylococcus aureus. Clin Microbiol Infect 2025;31:190–193. 10.1016/j.cmi.2024.10.022

54. Liu Q et al. Staphylococcus epidermidis Contributes to Healthy Maturation of the Nasal Microbiome by Stimulating Antimicrobial Peptide Production. Cell Host Microbe 2020;27:68–78.e5. 10.1016/j.chom.2019.11.003

55. Adolf LA, Heilbronner S. Nutritional Interactions between Bacterial Species Colonising the Human Nasal Cavity: Current Knowledge and Future Prospects. Metabolites 2022;12:489. 10.3390/metabo12060489

56. Rosenstein R et al. The Staphylococcus aureus-antagonizing human nasal commensal Staphylococcus lugdunensis depends on siderophore piracy. Microbiome 2024;12:213. 10.1186/s40168-024-01913-x

57. Bongers T. The Maturity Index, the evolution of nematode life history traits, adaptive radiation and cp-scaling. Plant Soil 1999;212:13–22.

58. Wang Y et al. Recovery of bacterial network complexity and stability after simulated extreme rainfall is mediated by K−/r-strategy dominance. Appl Soil Ecol 2024;203:105657. 10.1016/j.apsoil.2024.105657

59. Zheng W et al. The response patterns of r- and K-strategist bacteria to long-term organic and inorganic fertilization regimes within the microbial food web are closely linked to rice production. Sci Total Environ 2024;942:173681. 10.1016/j.scitotenv.2024.173681

60. Bushman SD, Skaar EP. The exploitation of nutrient metals by bacteria for survival and infection in the gut. PLOS Pathog 2025;21:e1013580. 10.1371/journal.ppat.1013580

61. Lucas SK et al. Anaerobic Microbiota Derived from the Upper Airways Impact Staphylococcus aureus Physiology. Infect Immun 2021;89:e00153–21. 10.1128/IAI.00153-21

62. Bonillo-Lopez L et al. Porcine nasal organoids to model interactions between the swine nasal microbiota and the host. Microbiome 2025;13:131. 10.1186/s40168-025-02088-9

63. Vallespir Lowery N, Ursell T. Structured environments fundamentally alter dynamics and stability of ecological communities. Proc Natl Acad Sci 2019;116:379– 388. 10.1073/pnas.1811887116

64. Junkins EN et al. Environmental structure impacts microbial composition and secondary metabolism. ISME Commun 2022;2:15. 10.1038/s43705-022-00097-5

