## Supplemental Figures for "Community composition drives metabolic competition and *Staphylococcus aureus* colonization resistance in synthetic nasal communities"

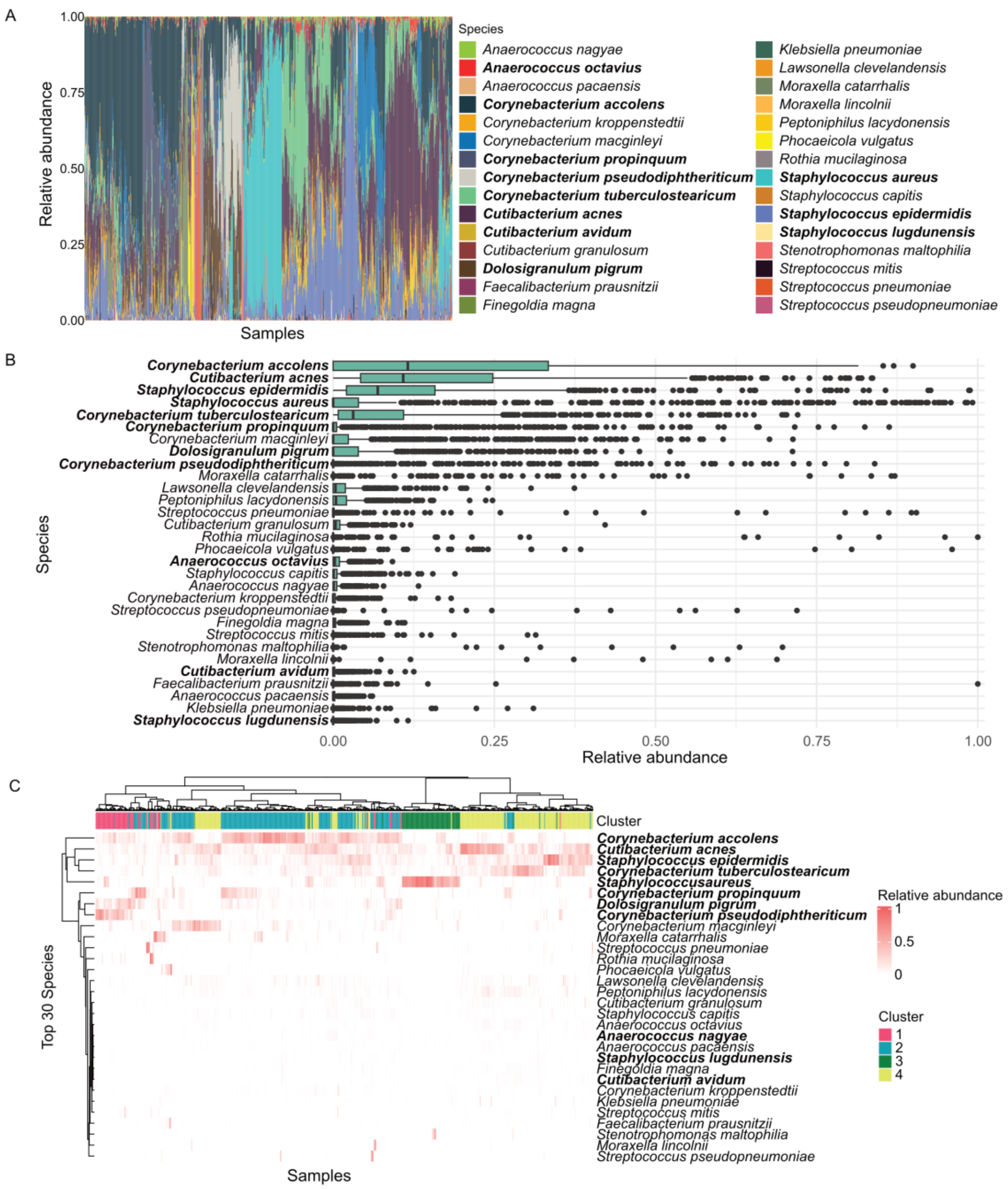


**Supplementary Figure 1. Human Microbiome Project nasal cavity data analysis and strain selection.** (A) Relative abundance barplots for 974 Human Microbiome Project nasal cavity samples. Relative abundances were calculated from 16S rRNA gene sequencing data. Only the 30 most abundant species across all samples are shown. Samples are ordered by compositional similarity using clustering of relative abundance community composition. The ten species selected for SynCom construction are highlighted in bold. (B) Boxplots showing the distribution of relative abundance for each of the top 30 species across all nasal samples. Each dot represents one sample, illustrating variability in each species abundance across all samples. Species are ordered by mean relative abundance. The ten species selected for SynCom construction are highlighted in bold. (C) Heatmap of relative abundances for the top 30 species across all samples. Samples were clustered based on Bray–Curtis dissimilarities of relative-abundance profiles. The optimal number of sample clusters (4) was chosen using the silhouette method and cluster membership is indicated in the top annotation. Columns are ordered according to hierarchical clustering with Ward's method on the same Bray–Curtis distance matrix. The ten species selected for SynCom construction are highlighted in bold.

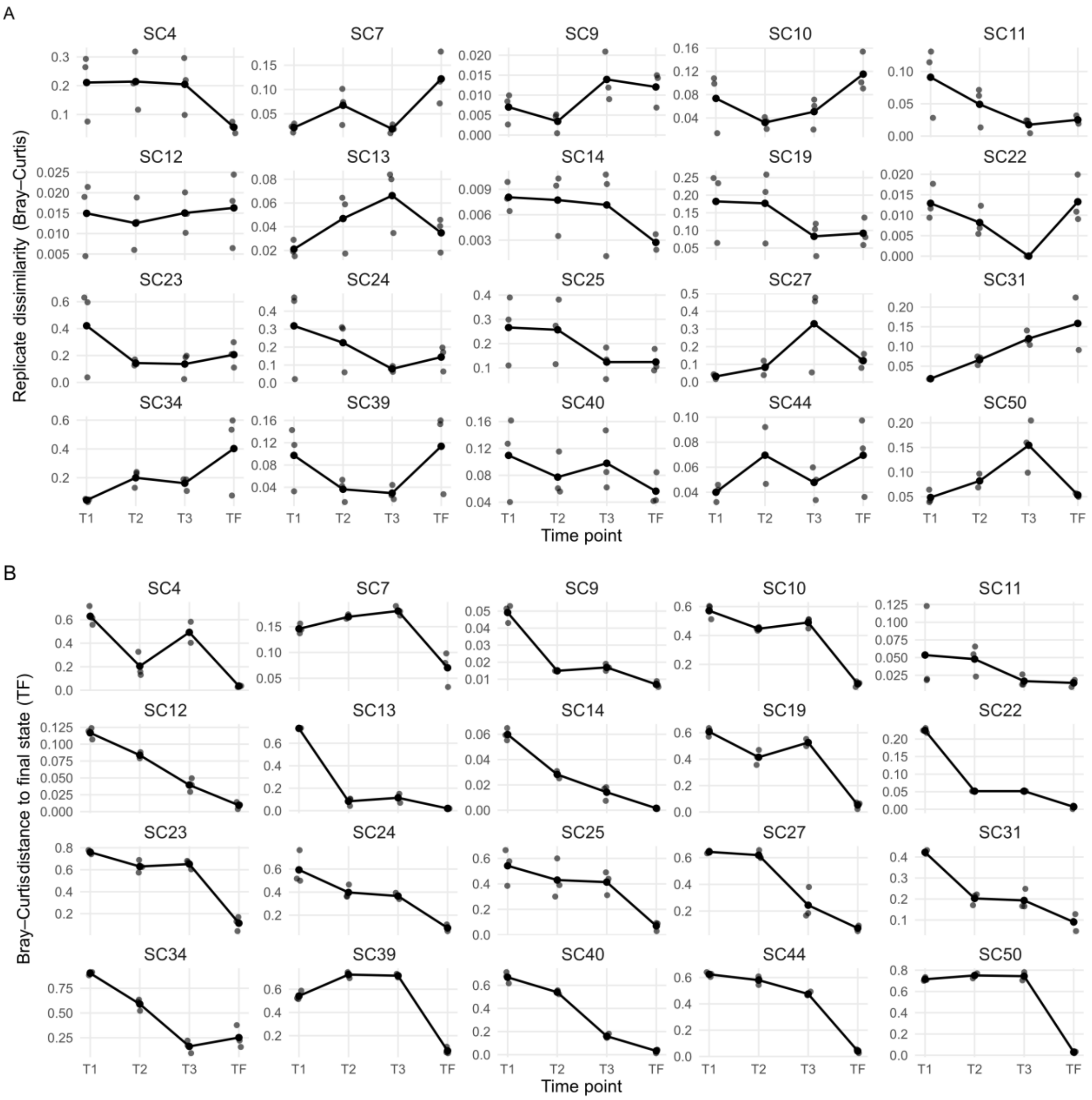


**Supplementary Figure 2. Replicate similarity and compositional stabilization across time.** (A) Similarity of biological replicates across all SynComs and time points. For each SynCom and time point, Bray–Curtis distances were computed between all pairs of replicates using relative abundance. Each point represents the mean pairwise distance among replicates at a given time point, with lower values indicating higher similarity. Panels correspond to individual SynComs. (B) Compositional stabilization trajectories for each SynCom. For every sample, Bray–Curtis distances to the corresponding final composition of each SynCom (i.e. the last time point) were calculated. These distances quantify how far each intermediate time point is from the final stabilized state. Decreasing distances across time reflect convergence of the replicate communities toward their final composition.

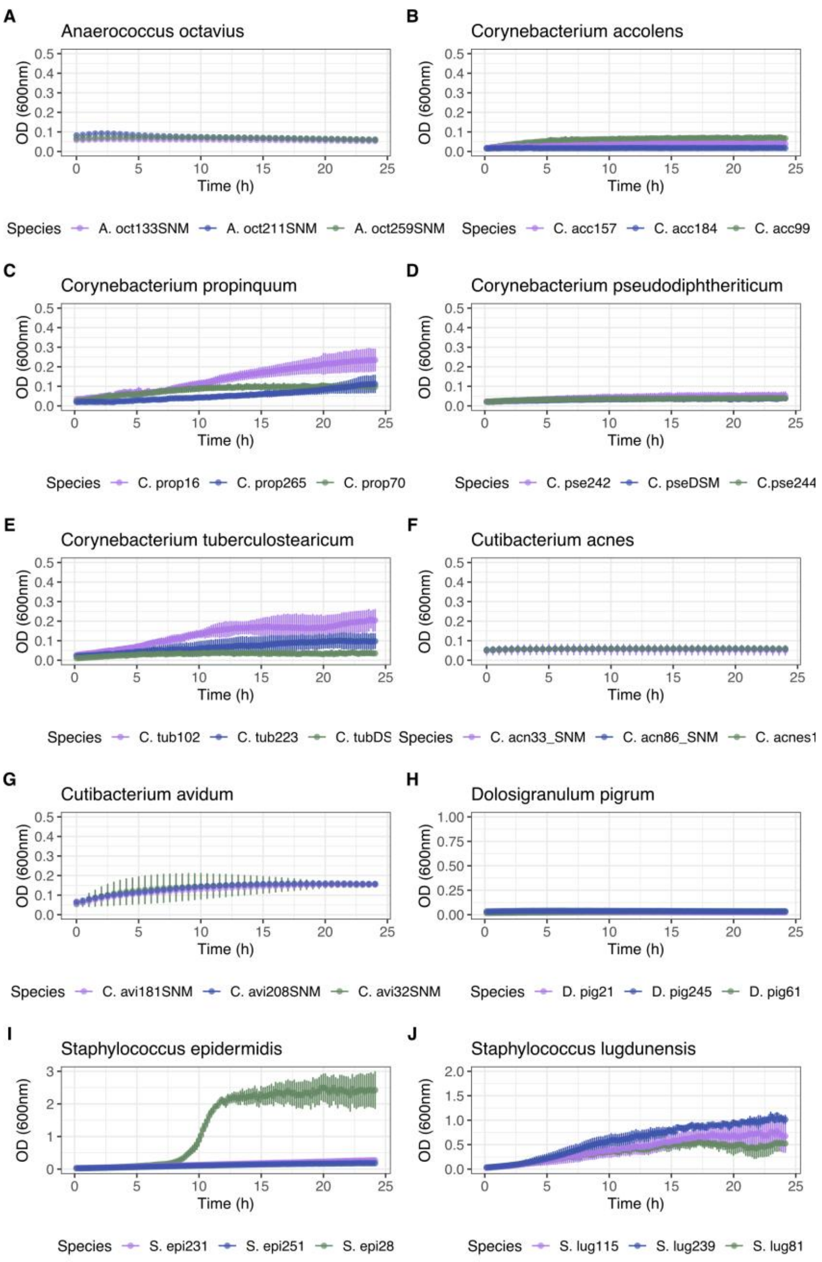


**Supplementary Figure 3. Growth curves of the 10 species (3 strains per species) in SNM3.** Each panel corresponds to one species and its 3 strains. Points show the mean optical density (OD) of four biological replicates per strain, measured every 10 minutes over 24 hours.

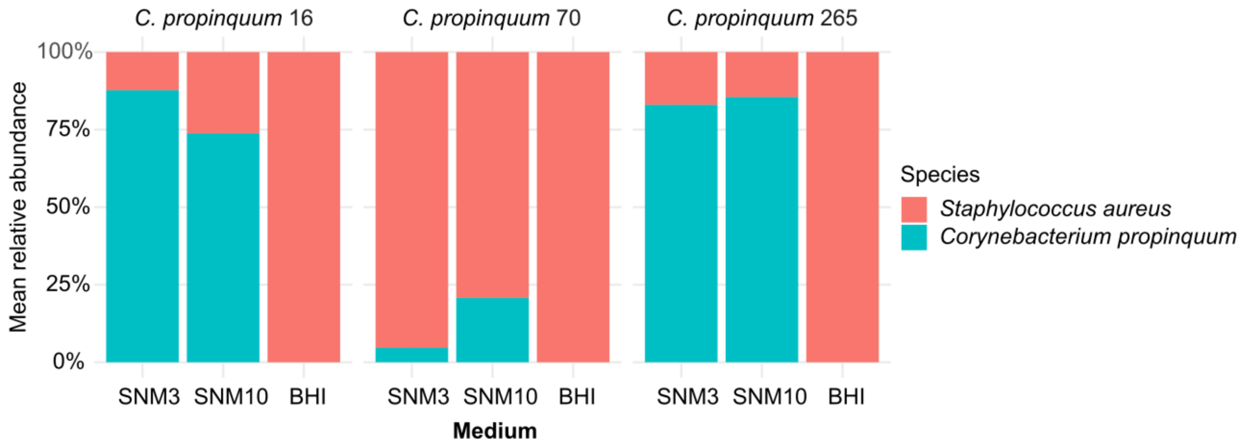


**Supplementary Figure 4. Coculture outcomes in SNM3, SNM10, and BHI.** Relative abundance of coculture experiments between the three *C. propinquum* strains used in the SynCom experiments and *S. aureus* USA300, the strain used to test colonization resistance. Each panel corresponds to one *C. propinquum* strain, and barplots display the mean relative abundance across three biological replicates.
